# Targeting TrkB with a brain-derived neurotrophic factor mimetic promotes myelin repair in the brain

**DOI:** 10.1101/268300

**Authors:** Jessica L Fletcher, Rhiannon J Wood, Jacqueline Nguyen, Eleanor ML Norman, Christine MK Jun, Alexa R Prawdiuk, Melissa Biemond, Huynh TH Nguyen, Susan E Northfield, Richard A Hughes, David G Gonsalvez, Junhua Xiao, Simon S Murray

**Affiliations:** Department of Anatomy and Neuroscience, School of Biomedical Science, Faculty of Medicine, Dentistry and Health Sciences, The University of Melbourne, Parkville, 3052, VIC, AUSTRALIA; Department of Pharmacology and Therapeutics School of Biomedical Science, Faculty of Medicine, Dentistry and Health Sciences, The University of Melbourne, Parkville, 3052, VIC, AUSTRALIA

**Keywords:** remyelination, brain-derived neurotrophic factor, oligodendrocyte, TrkB

## Abstract

Methods to promote myelin regeneration in response to central myelin loss are essential to prevent the progression of clinical disability in demyelinating diseases. The neurotrophin brain-derived neurotrophic factor (BDNF) is known to promote myelination during development *via* oligodendrocyte expressed TrkB receptors. Here, we use a structural mimetic of BDNF to promote myelin regeneration in a preclinical mouse model of central demyelination. We show that selective targeting of TrkB with the BDNF-mimetic enhances remyelination, increasing oligodendrocyte differentiation, the frequency of myelinated axons, and myelin sheath thickness after a demyelinating insult. Treatment with exogenous BDNF exerted an attenuated effect, increasing myelin sheath thickness only. Further, following conditional deletion of TrkB from pre-myelinating oligodendrocytes, we show the effects of the BDNF-mimetic on oligodendrocyte differentiation and remyelination are lost, indicating these are dependent on oligodendrocyte expression of TrkB. Overall, these studies demonstrate that targeting oligodendrocyte TrkB promotes *in vivo* remyelination in the brain.

## Introduction

Innate myelin regeneration is often incomplete in central nervous system (CNS) demyelinating diseases such as multiple sclerosis (MS). This leaves axons exposed, resulting in conduction deficits, loss of metabolic and trophic support to axons, and contributes to secondary irreversible axonal damage that ultimately drives the progressive clinical disability in patients (Lucchinetti *et al.*, 1999; Chang *et al.*, 2002; Albert *et al.*, 2007). Current commercially available immunomodulatory therapies for MS effectively decrease the frequency of relapses, but do not directly stimulate remyelination (Stangel *et al.*, 2017). As such there is a compelling need to complement these therapies with new strategies to promote myelin regeneration. The neurotrophin brain-derived neurotrophic factor (BDNF) enhances CNS myelination, acting through oligodendrocyte-expressed TrkB receptors (Du *et al.*, 2003; Xiao *et al.*, 2011). Despite the relative potency with which BDNF enhances myelination, its promiscuity to both p75^NTR^ and TrkB receptors, brief half-life (Poduslo and Curran, 1996), large molecular size and relatively poor ability to cross the blood brain barrier make it a poor therapeutic candidate, likely contributing to its failure in clinical trials for neurodegenerative disease (The BDNF Study Group, 1999; Beck *et al.*, 2005).

To overcome these limitations, a range of small molecular weight neurotrophin receptor agonists have been developed (Longo and Massa, 2013). This includes tricyclic-dimeric peptide-6 (TDP6), a peptide mimetic structurally based on the loop-2 region of the BDNF homodimer that interacts with TrkB (O’Leary and Hughes, 2003). When BDNF binds and activates TrkB, it triggers autophosphorylation of the intracellular domain, recruitment of cytosolic adaptor proteins and activation of many intracellular signaling cascades, including PI3K/Akt and MAPK/Erk (Chao, 2003). We have previously shown that TDP6, like BDNF, enhances oligodendrocyte myelination *in vitro*, and does this through phosphorylation of oligodendrocyte expressed TrkB receptors (Wong *et al.*, 2014).

There is a clinical need to develop remyelinating therapies capable of complementing existing immunomodulatory treatments for MS. Here we test whether oligodendrocyte expressed TrkB receptors are a rational target to achieve this, by assessing whether TDP6 promotes myelin repair *in vivo*. We show infusion of TDP6, but not BDNF, into the CNS increased oligodendrocyte differentiation, the frequency of myelinated axons and myelin sheath thickness in the cuprizone model of toxic demyelination. Importantly, we demonstrate that this effect is driven by phosphorylation of oligodendrocyte expressed TrkB receptors *in vivo*. These data suggest selective targeting of TrkB is a rational approach to promote myelin repair *in vivo.*

## Results

### BDNF, and its structural mimetic TDP6, enhance myelin repair following cuprizone-induced demyelination

Having previously shown that BDNF and TDP6 enhance oligodendrocyte myelination *in vitro* (Wong *et al.*, 2014), here we tested the capacity of BDNF and TDP6 to promote remyelination *in vivo*. To do this, demyelination was induced in 8-10 week-old female C57BL/6 mice with 0.2% cuprizone in normal chow for 6 weeks (Fig 1A). Efficacy of cuprizone-induced demyelination was confirmed through electron microscopy (EM) and immunostaining for myelin basic protein (MBP) in the brains of mice collected at the end of the 6-week cuprizone challenge (minimum *n*=2/cohort). These mice demonstrated demyelination, with loss of myelin from axons identified by EM and severe reduction in the level of MBP expression within the corpus callosum (Fig. S1). Remaining mice received 4μm BDNF, 40μM TDP6 or artificial cerebrospinal fluid vehicle delivered at a rate of 0.5μL/hr by intracerebroventricular osmotic pump for 7 days to the right lateral ventricle. Dose was chosen based on our previous *in vitro* data (Wong *et al.*, 2014). After 7 days infusion, immunostaining (Fig. 1B) revealed that BDNF and TDP6 significantly increased the area of the contralateral caudal corpus callosum positive for MBP (Fig 1C; ****p<0.001), indicative of greater remyelination. EM analysis of myelin ultrastructure (Fig. 1B) revealed that in the caudal corpus callosum of mice treated with TDP6, the proportion of myelinated axons was significantly increased compared to vehicle controls (Fig. 1D, *p<0.018). In contrast, in those treated with BDNF, the proportion of myelinated axons was similar to those that only received the vehicle (Fig. 1D). Assessment of myelin sheath thickness by g-ratio revealed that TDP6 treatment significantly reduced the mean g-ratio compared to the vehicle (Fig. 1E, *p=0.019) indicating thicker myelin sheaths. Linear regression indicated that BDNF also significantly increased myelin sheath thickness (Fig. 1F, *p=0.04) compared to the vehicle, but that TDP6 had a more pronounced effect with significantly thicker myelin than both BDNF (Fig. 1F, *p=0.02) and vehicle treatment (Fig. 1F, **p=0.014). There was no change in the frequency distribution of myelinated axons based on axon diameter (Fig. 1G). Overall, these findings indicate TDP6 significantly enhances remyelination *in vivo*.

**Fig. 1:**
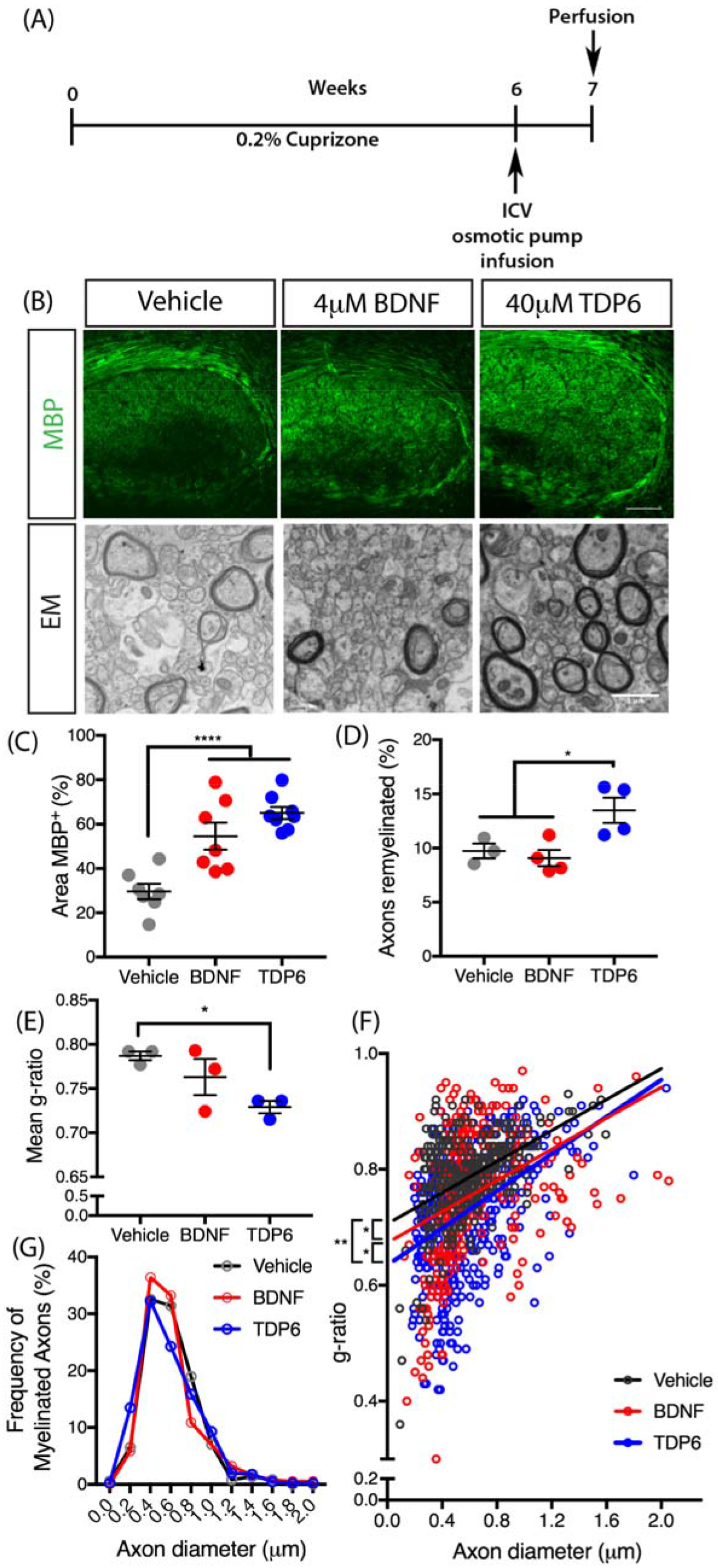
B DNF and TDP6 treatment enhance remyelination following cuprizone-induced demyelination. **(A)** Schematic of experimental procedures. **(B)** Representative micrographs of myelin basic protein (MBP) immunostaining, and electron micrographs (EM) of myelinated and unmyelinated axons in the caudal corpus callosumof vehicle, BDNF and TDP6 treated mice respectively. MBP micrographs: scale bar=20μm. EM: scale bar=1μm **(C)** Percentage area positive for MBP immunostaining was significantly enhanced in the caudal corpus callosum of mice treated with 4μM BDNF and 4μMM TDP6 for 7 days after 6-weeks cuprizone demyelination, compared to the vehicle treated mice (****p<0.0001, *n*=7-9/group). ***(D)***Percentage of myelinated axons was significantly increased with TDP6, but not BDNF, compared to vehicle treated controls (*p=0.018, *n*=3-4/group). **(E)** Mean g-ratio was significantly decreasedwith TDP6 treatment compared to vehicle controls, indicative of thicker myelin sheaths(*p=0.019,*n*=3-4/group). For *(C-E)* 1-way ANOVA with Tukey’s post-hoc multiple comparisons. Mean ±SEM plotted. **(F)** Scatter-plot of g-ratio against axon diameter. Linear regression analysis revealed a significant decrease in the y-intercepts between BDNF and TDP6treated mice compared to vehicle treated controls, and between TDP6 compared to BDNF treatment (*p=0.0042), but no significant change in slope (p=0.45). **(G)** Frequency distribution plot of myelinated axon diameters indicatingno change with treatment in the frequency of myelinated axons based on size (p=0.43, χ^2^ distribution test). For *(D-G)* Minimum 150axons/animal, *n*=3/group.

### Treatment with TDP6, but not BDNF, increases oligodendrocyte density and differentiation

To identify whether BDNF or TDP6 treatment exerted an effect on oligodendroglial sub-populations, we next examined the density of total Olig2^+^ oligodendroglia, Olig2^+^PDGFRα^+^ oligodendrocyte progenitor cells (OPCs) and post-mitotic Olig2^+^CC1^+^ oligodendrocytes in the caudal corpus callosum using immunohistochemistry (Fig. 2A). The density of Olig2^+^ oligodendroglia was significantly increased with TDP6 treatment, above that seen with BDNF or vehicle treatment (Fig. 2B, *p=0.024). Examination of the contribution of OPCs to this increase in oligodendroglia, revealed there was no significant change in the density of Olig2^+^PDGFRα^+^ OPCs in mice treated with TDP6 compared to those treated with BDNF or the vehicle (Fig. 2C, p=0.06). In contrast, the density of Olig2^+^CC1^+^ oligodendrocytes in TDP6 treated mice was significantly increased compared to treatment with BDNF and the vehicle (Fig. 2D, **p=0.0051). This increased density of oligodendrocytes, but not OPCs, suggests TDP6 may increase overall oligodendrocyte differentiation and/or their survival during remyelination. Together, the increase in oligodendrocyte density is consistent with the increased proportion of remyelinated axons in mice treated with TDP6, but not BDNF.

**Fig. 2:**
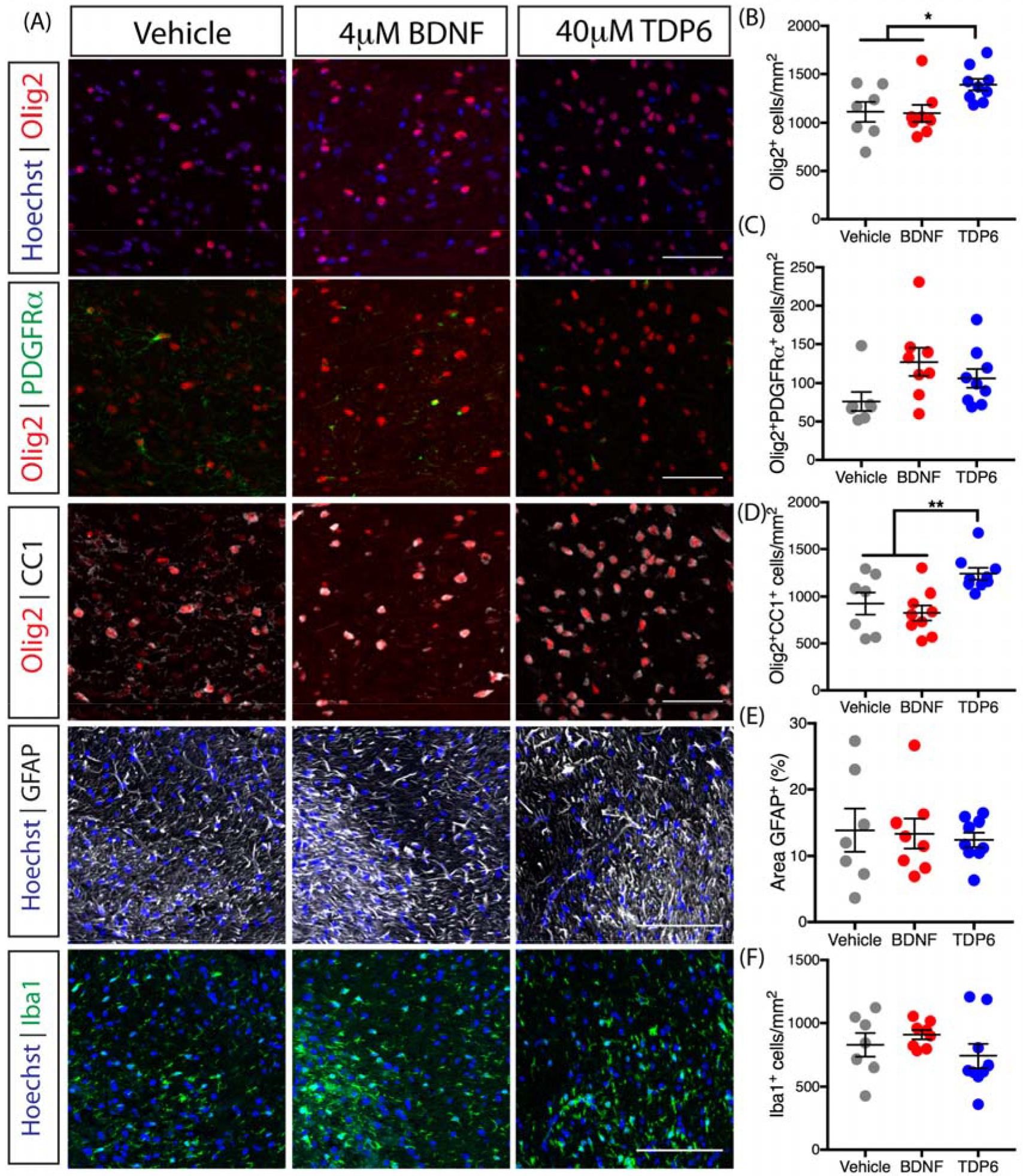
Treatment with TDP6, but not BDNF, enhances oligodendrocyte density and differentiation during remyelination. **(A)** Representative micrographs of immunostaining in the caudal region of the corpus callosum for total Olig2^+^ oligodendrocytes, Olig2^+^PDGFRα^+^ OPCs, and Olig2^+^CC1^+^ oligodendrocytes (scale bar=20μm), as well as GFAP^+^ astrocytosis and Iba1^+^ microglia (scale bar=100μm). **(B)** Density of total Olig2^+^ oligodendrocytes was significantly increased in TDP6 treated mice compared to treatment with BDNF or vehicle (*p=0.024). **(C)** Density of Olig2^+^PDGFRα^+^ OPCs was unchanged across treatments (p=0.06). **(D)** Density of Olig2^+^CC1^+^ oligodendrocytes significantly increased with TDP6 treatment compared to BDNF and vehicle treatments (*p=0.0051). **(E)** Percentage area of GFAP^+^ astrocytes was unchanged across treatments (p=0.89). **(F)** Density of Iba1^+^ microglia was unchanged across treatments (p=0.34). For *(B-F) n*=7-9/group, 1-way ANOVA with Tukey’s post-hoc multiple comparisons. Mean ±SEM plotted.

To determine if BDNF or TDP6 treatment altered gliosis, the contralateral caudal corpus callosum was immunolabelled with GFAP for astrocytosis and Iba1 for microgliosis (Fig. 2A). Astrocytosis persisted with both BDNF and TDP6 treatments, and the level of GFAP staining was unchanged across the three groups (Fig. 2E, p=0.34). Similarly, the density of Iba1^+^ microglia was unchanged across treatments (Fig. 2F, p=0.34). This suggests treatment with either TDP6 or BDNF exerts no reductive effects on neuroinflammatory cell populations, indicating their effect on myelin repair is not secondary to an anti-inflammatory effect.

### TDP6 and BDNF increase TrkB phosphorylation in oligodendrocytes during remyelination

As a structural-mimetic of the loop-2 region of the BDNF homodimer, TDP6 is designed to selectively interact with and initiate phosphorylation of TrkB receptors (O’Leary and Hughes, 2003). Indeed, we have previously shown that TDP6 phosphorylates oligodendrocyte-expressed TrkB receptors and promotes myelination *in vitro* (Wong *et al.*, 2014). To examine if BDNF and TDP6 treatment successfully stimulated TrkB phosphorylation in oligodendrocytes *in vivo*, the contralateral caudal corpus callosum was co-immunolabelled with antibodies directed against the phosphorylated serine 478 of (pTrkB^S478^), and the oligodendroglial markers CC1 and PDGFRα) (Fig. 3A). This revealed that the proportion of total pTrkB^S478+^ cells in the corpus callosum was significantly increased following TDP6 infusion, but not BDNF, compared to the vehicle (Fig. 3B,**p=0.0047). Assessing the proportion of pTrkB^S478+^ cells that were PDGFRα^+^ OPCs, indicated a trend towards increased pTrkB^S478+^ OPCs in mice treated with TDP6 (Fig. 3C, p=0.12). The proportion of CC1-pTrkB^S478^ double-positive cells in the corpus callosum was significantly increased in mice treated with BDNF and TDP6 (Fig. 3D, ****p<0.0001).These results demonstrate that both BDNF and TDP6 successfully reached their cellular targets following intracerebroventricular delivery, and were capable of phosphorylating TrkB on oligodendrocytes.

**Fig. 3:**
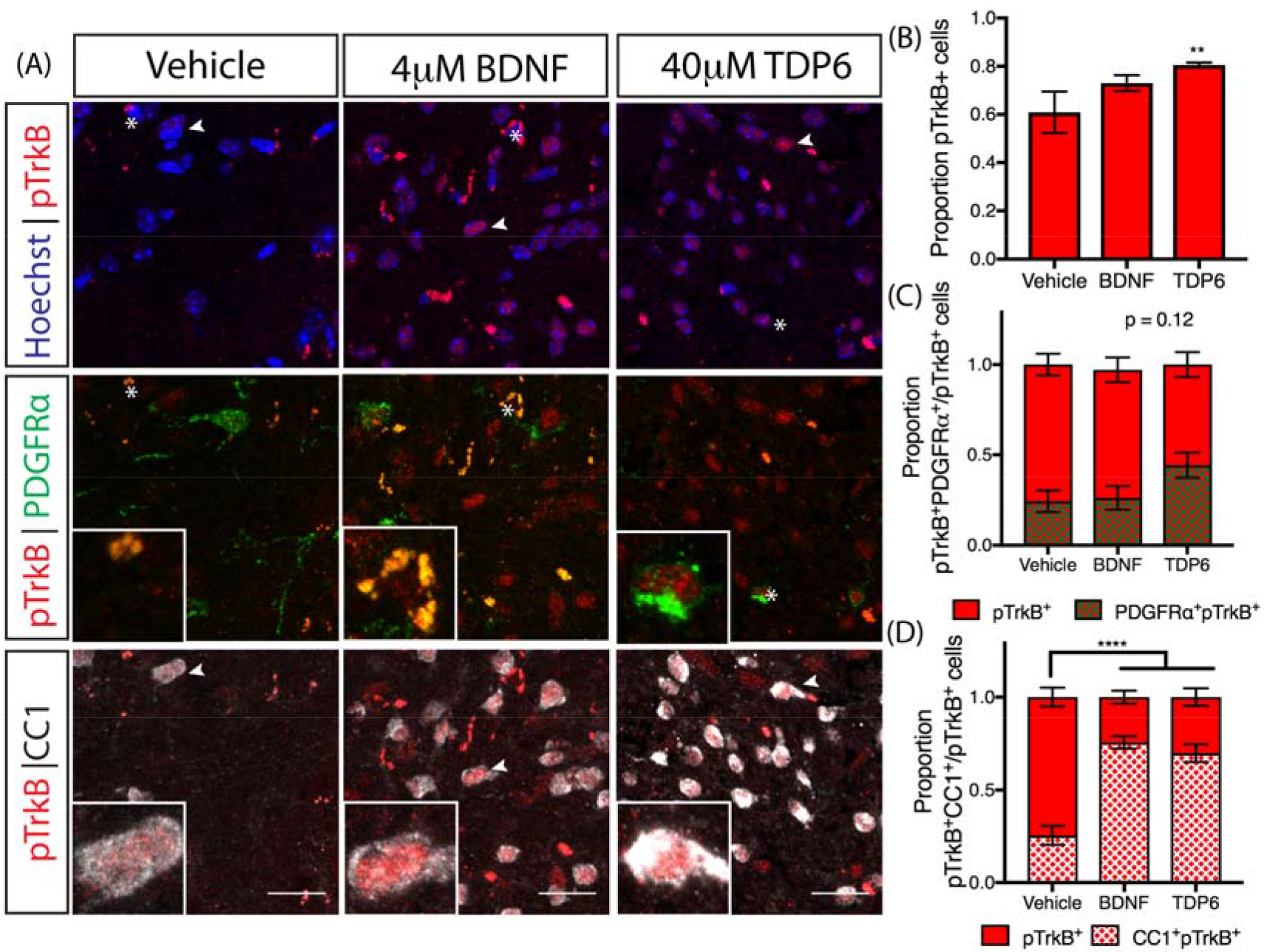
Treatment with TDP6 increased TrkB phosphorylation in the remyelinating corpus callosum. **(A)** Representative micrographs of pTrkB^S478^ immunostaining in the caudal region of the corpus callosum. Scale bar=202m. asterisk: PDGFRα^+^ cell featured in inset, arrowhead: CC1^+^ cell featured in inset. **(B)** Proportion of pTrkB^S478+^ cells in the caudal corpus callosum was increased with TDP6 treatment compared to BDNF and vehicle controls (**p=0.0047). **(C)** Proportion of pTrkB^S478+^ cells that were PDGFRα^+^ OPCs tended towards an increase with TDP6 treatment (p=0.12). **(D)** Proportion of pTrkB^S478+^ cells that were CC1^+^ oligodendrocytes significantly increased with both BDNF and TDP6 treatment. For *(B-D)* n=7-9/group, 1-way ANOVA with Tukey’s post-hoc multiple comparisons. Mean ± SEM plotted.

### Adult myelination and demyelination is unaltered by deletion of oligodendroglial TrkB

To test if the pro-remyelinating effect of TDP6 required TrkB expression on oligodendrocytes, we generated mice with an oligodendrocyte specific deletion of TrkB (Lulkart *et al.*, 2005) driven by the CNPase promoter (Lappe-Siefke *et al.*, 2003) (CNPaseCre^+/−^ TrkB^fl/fl^ mice). These mice had approximately 3-fold reduction in Olig2^+^TrkB^+^ cells in the caudal region of the corpus callosum at 14-16 weeks of age (Fig. S2A, quantified in Fig. S2B).

To assess the effect of TrkB deletion from CNPase-expressing cells on oligodendrocyte density and myelin, we first compared unchallenged conditional knockout and wildtype littermate mice, then compared the effect of cuprizone-induced demyelination on the two genotypes. Female CNPaseCre^−/−^ TrkB^fl/fl^ and CNPaseCre^+/−^ TrkB^fl/fl^ mice were unchallenged, or fed 0.2% cuprizone in normal chow for 6 weeks from 8-10 weeks of age, sacrificed and the caudal corpus callosum immunohistochemically analyzed (Fig. S2C). In unchallenged mice, similar levels of MBP staining were observed regardless of genotype, and both genotypes demyelinated to a similar extent following cuprizone (Fig. S2C, quantitated in Fig.S2D). Similarly, the density of Olig2^+^ oligodendroglia (Fig. S2E), Olig2^+^PDGFRpα^+^ OPCs (Fig. S2F) and Olig2^+^CC1^+^ oligodendrocytes (Fig. S2G) was similar in unchallenged mice, with cuprizone exerting no differences between genotypes. Assessment of astrogliosis (GFAP staining, Fig. S2H quantitated in Fig. S2I) and microgliosis (Iba1 staining, Fig. S2H quantitated in Fig. S2J) also exhibited no differences between genotype in either unchallenged or cuprizone challenged conditions. These data indicate that oligodendroglial TrkB does not exert an essential role in oligodendroglial or myelin maintenance in the adult, or following cuprizone-induced demyelination

### TDP6-enhanced oligodendrocyte differentiation is dependent on oligodendroglial TrkB

To determine if the pro-remyelinating effects of TDP6 required oligodendrocyte expression of TrkB, we repeated the cuprizone experiments and 7-day osmotic pump infusions with either TDP6 (*n*=4) or vehicle (*n*=3) in CNPaseCre^+/−^ x TrkB^fl/fl^ conditional knockout mice. Immunohistochemistry for MBP (Fig. 4A) revealed the pro-remyelinating effect exerted by TDP6 in wildtype mice was lost, with both vehicle and TDP6 treated conditional knockout mice exhibiting the same level of MBP staining (Fig. 4B, p=0.45). Further, the density of total Olig2^+^ oligodendroglia was also unchanged following TDP6 treatment (Fig. 4C, p=0.55) and there was no change in the density of Olig2^+^PDGFRpα^+^ OPCs or Olig2^+^CC1^+^ oligodendrocytes between treatment groups in the conditional knockout mice (Fig. 5D-E, p=0.80 and p=0.17 respectively). These data suggest the effect TDP6 exerts to promote remyelination and increase oligodendrocyte density is dependent on oligodendrocyte expressed TrkB.

**Fig. 4:**
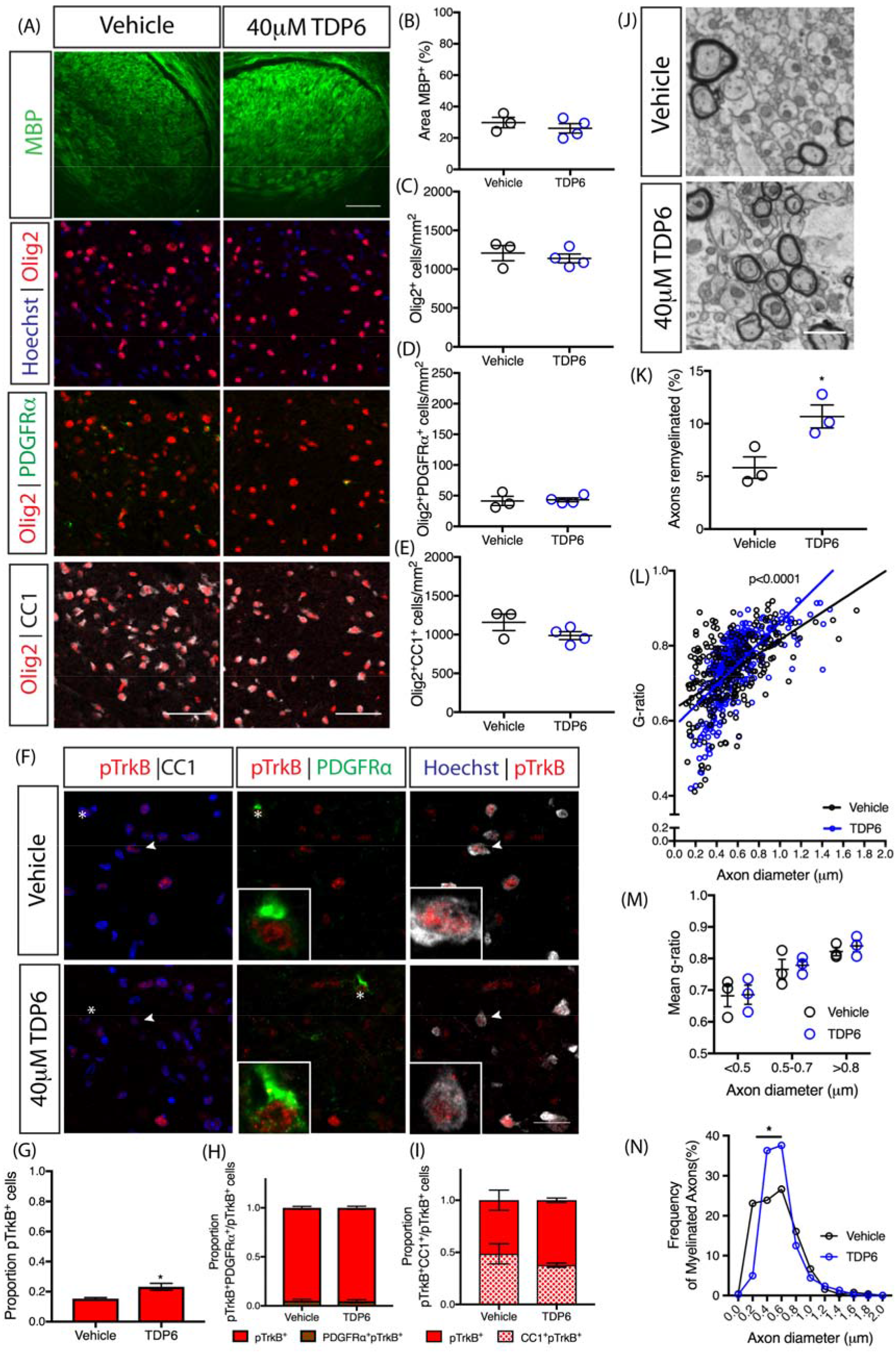
Enhanced oligodendrocyte differentiation and increases in myelin sheath thickness by TDP6 during remyelination require oligodendrocyte TrkB. **(A)** Representative micrographs of MBP immunostaining and oligodendrocyte lineage markers in the caudal corpus callosum of CNPasCre^+/−^ TrkB^fl/fl^ cuprizone-demyelinated mice treated with vehicle or TDP6. Scale bar=20μm. **(B)** Percentage area of MBP immunostaining (p=0.45), density of **(C)** Olig2^+^ cells (p=0.55) **(D)** Olig2^+^PDGFRα^+^ OPCs (p=0.80) and **(E)** Olig2^+^CC1^+^ oligodendrocytes (p=0.17) were unchanged between vehicle and TDP6 treated TrkB conditional knockout mice. Unpaired Student’s t-test, n=3-4/group. **(F)** Representative micrographs of pTrkB^S478^ immunostaining in the caudal corpus callosum of CNPasCre^+/−^ TrkB^fl/fl^ cuprizone-demyelinated mice treated with vehicle or TDP6. Scale bar=20μm. **(G)** Proportion of pTrkB^S478+^ cells were significantly increased in TDP6 treated conditional knockout mice (p=0.037), but **(H)** the proportion of pTrkB^S478+^ PDGFRα^+^ OPCs were unchanged (p=0.80) as were **(I)** the proportion of pTrkB^S478+^ CC1^+^ oligodendrocytes (p=0.25). Unpaired student’s t-test, n=3-4/group. **(J)** Electron micrographs of the caudal corpus callosum of CNPaseCre^+/−^ TrkB^fl/fl^ cuprizone-demyelinated mice treated with vehicle or TDP6. Scale bar=1μm. **(K)** Proportion of myelinated axons was significantly increased in TrkB conditional knockout mice that received TDP6 compared to vehicle controls (*p=0.0314). **(L)** Scatter plot of g-ratio against axon diameter. Linear regression analysis revealed a significant increase in slope between CNPaseCre^+/−^ TrkB^fl/fl^ mice treated with TDP6 compared to vehicle (****p<0.0001). **(M)** Mean g-ratio categorized by axonal diameter revealed no significant change between treatment groups, indicative of no change in myelin sheath thickness with TDP6 treatment in TrkB conditional knockout mice (p=0.959), n=3/group 2-way ANOVA. **(N)** Frequency plot of myelinated axon diameters revealing a significant increase in the frequency of axons myelinated ranging from 0.3-0.6om diameter in CNPaseCre^+/−^ TrkB^fl/fl^ mice treated with TDP6 (p=0.0032, χ^2^ distribution test). For (H-J) minimum 100 axons/animal, n=3/group.

Successful delivery of TDP6 was confirmed by immunohistochemistry for pTrkB^S478^ (Fig. 4F) which revealed that the proportion of total pTrkB^S478+^ cells in the corpus callosum significantly increased in conditional knockout mice that received TDP6, compared to those receiving the vehicle (Fig. 4G, *p=0.037). Consistent with reduced levels of TrkB expression in the CNPaseCre^+/−^ TrkB^fl/fl^ mice, there was a low proportion of pTrkB^S478+^ PDGFRα^+^ OPCs(Fig. 4H) and pTrkB^S478+^ CC1^+^ oligodendrocytes (Fig. 4I), and these were unchanged by treatment with TDP6 (p=0.80 and p=0.25, respectively). Delivery of TDP6 was further verified by pooling residual TDP6-solution remaining in the osmotic pump reservoir at the end of the treatment period from the conditional knockout mice, and run through reverse-phase HPLC. This demonstrated that TDP6 was stable within the reservoir throughout the treatment period (Fig. S3A-D).

Despite no change in MBP staining, EM analysis (Fig. 4J) revealed that the proportion of axons remyelinated in the caudal corpus callosum was significantly increased with TDP6 treatment in conditional knockout mice (Fig. 4K, *p=0.031). However, the mean g-ratio was unchanged (Vehicle: 0.74±0.02; TDP6: 0.75±0.02, *n*=3/group, p=0.76), indicating that while TDP6 increased the proportion of myelinated axons, it exerted no effect on myelin sheath thickness in the conditional knockout mice. Linear regression analysis revealed that compared to vehicle treatment, TDP6 in the conditional knockout mice resulted in a significant increase in slope (Fig. 4L, p<0.0001), suggestive of an altered relationship between myelin profiles and axonal diameter. To dissect the driver of this effect, g-ratios were categorized based on axon diameter. This revealed that there was no change in myelin sheath thickness across axons of different diameter (Fig. 4M), suggesting the change in slope was likely driven by an increase in the number of axons myelinated in each size category between treatments. Consistent with this, analysis of the frequency distribution of myelinated axons based on their diameter demonstrated that there were significantly more axons between 0.3-0.6μm range that were myelinated in TDP6 treated conditional knockout mice (Fig. 4N, *p=0.0032). This suggests that TDP6 may exert a small oligodendroglial-TrkB independent effect to initiate myelin ensheathment for a selective subset of axons, but not increase myelin sheath thickness.

## Discussion

In the present study, our data show selective targeting of TrkB receptors with the BDNF mimetic, TDP6 in the cuprizone-demyelinated brain enhanced remyelination. TDP6 infusion increased the density of Olig2^+^CC1^+^ oligodendrocytes, proportion of remyelinated axons, and myelin sheath thickness following 7-days recovery. Infusion with BDNF, the neurotrophin from which TDP6 is derived, demonstrated an attenuated response, only increasing myelin sheath thickness. Importantly, the effects of TDP6 required expression of oligodendrocyte TrkB receptors. Intriguingly,following conditional deletion of TrkB from oligodendroglia, TDP6 retained some capacity to increase the proportion of remyelinated axons, but not myelin sheath thickness. This suggests that TrkB activation may exert segregated effects, whereby oligodendrocyte expression of TrkB increases oligodendrocyte differentiation and myelin sheath thickness, whereas an alternate source of TrkB, potentially neuronal, initiates myelin ensheathment, effectively increasing the frequency of remyelinated axons following TDP6 treatment.

The therapeutic potential of BDNF has been tested in numerous intervention studies and clinical trials for neurologic conditions over the past several decades (McTigue *et al.*, 1998; The BDNF Study Group, 1999; Beck *et al.*, 2005; Makar *et al.*, 2009; Fulmer *et al.*, 2014; Ramos-Cejudo *et al.*, 2015; Gonsalvez *et al.*, 2017). In demyelinating diseases including MS, direct infusion (Ramos-Cejudo *et al.*, 2015), cell-based gene therapy to overexpress BDNF (McTigue *et al.*, 1998; Makar *et al.*, 2009), and indirect modulation of endogenous BDNF secretion (Fulmer *et al.*, 2014) in animal models have shown promise to improve remyelination However, by and large, these studies utilized indirect measures of remyelination, such as OPC proliferation and increased expression of myelin proteins, MBP, MOG and PLP (McTigue *et al.*, 1998; Makar *et al.*, 2009; Fulmer *et al.*, 2014; Ramos-Cejudo *et al.*, 2015). We have demonstrated that while direct BDNF infusion to the demyelinated CNS increased MBP, and tended to increase OPC density, consistent with previous reports (McTigue *et al.*, 1998; Makar *et al.*, 2009; Fulmer *et al.*, 2014; Ramos-Cejudo *et al.*, 2015), it did not increase the proportion of myelinated axons, or oligodendrocyte differentiation. These abilities are essential to a viable remyelinating strategy (Murphy and Franklin, 2017), particularly for MS lesions where OPC differentiation appears arrested (Chang *et al.*, 2002). In contrast, by selectively targeting TrkB, the direct molecular mechanism used by BDNF to promote myelination in development (Du *et al.*, 2003; Xiao *et al.*, 2011; Wong *et al.*, 2013), we increased the density of Olig2^+^CC1^+^ oligodendrocytes and the proportion of myelinated axons. In this respect targeting oligodendroglial TrkB with TDP6 meets the criteria of a potential remyelinating therapeutic strategy.

It is well appreciated that BDNF promotes CNS myelination through activation of oligodendrocyte expressed TrkB (Du *et al.*, 2003; Xiao *et al.*, 2011; Wong *et al.*, 2013). We found that selective targeting of TrkB, with TDP6, led to greater TrkB phosphorylation in the corpus callosum, more mature oligodendrocytes, and higher frequency of remyelinated axons than exogenous BDNF. The precise reason why TDP6 elicited a significantly greater remyelinating effect than BDNF remains unclear. Deductive reasoning suggests potential differences in specificity, stability, dosage, or a combination of all three, may contribute to this discrepancy. The concentrations used were based on previous *in vitro* myelinating co-cultures where 25-fold more TDP6 was required to increase the number of MBp^+^ myelinated axonal segments to the same level as BDNF, and induce TrkB and Erk1/2 phosphorylation (Wong *et al.*, 2014). From this *in vitro* analysis, BDNF is substantially more potent than TDP6, suggesting it is unlikely the lower concentration of BDNF caused the attenuated response. The notoriously labile and highly charged nature of BDNF may contribute to its reduced efficacy *in vivo*. BDNF is known to have a very short half-life [7], while here we have shown that TDP6 can be retrieved from the osmotic pump reservoirs after 7-days *in vivo* (Fig. S3). Finally, the receptor specificity may also play a role. TDP6 selectively interacts with TrkB and has not shown any ability to interact with p75^NTR^ (O’Leary and Hughes, 2003; Xiao *et al.*, 2011; Wong *et al.*, 2014). It is possible that BDNF-p75^NTR^ interactions may have attenuated the BDNF response. Regardless, the attenuated remyelinating response to BDNF is consistent with reduced TrkB phosphorylation observed compared to TDP6. Importantly, compared to other BDNF-mimetics, we have shown using conditional knockout mice that the effects of TDP6 are predominantly dependent on expression of TrkB by the target cell *in vivo.* While other BDNF-mimetics have demonstrated benefits in various neurodegenerative conditions (Jiang *et al.*, 2013; Simmons *et al.*, 2013), this is the first time the functional effect of a neurotrophin mimetic has been directly identified to require its targeted receptor on a specific cell type. Overall, our results demonstrate the poor therapeutic potential of BDNF, and support the strategy to develop functional BDNF mimics as therapies for neurologic disease.

A feature consistent with both BDNF and TDP6 treatment was increased myelin sheath thickness, indicated by reduced g-ratios. Importantly, this is the first time this effect of exogenous BDNF has been demonstrated on myelin sheath *in vivo*. A role for BDNF in modulating myelin sheath thickness is not unexpected. BDNF^−/−^, BDNF^+/−^ and MBPCre^+/−^ x TrkB^fl/fl^ mice all demonstrate developmental hypomyelination with thinner myelin sheaths compared to wildtype mice (Cellerino *et al.*, 1997; Xiao *et al.*, 2011;Wong *et al.*, 2013). In addition, BDNF-TrkB signaling activates the MAPK/Erk pathway (Du *et al.*, 2006; Xiao *et al.*, 2012), which when activated in oligodendrocytes increases myelin sheath thickness (Ishii *et al.*, 2012, 2016; Fyffe-Maricich *et al.*, 2013). The effect that BDNF and TDP6 exert on oligodendrocytes to increase myelin sheath thickness requires TrkB expression, as there was no change in g-ratio when TDP6 was delivered during remyelination in the TrkB conditional knockout mice.

Similarly, the effect TDP6 exerted to increase oligodendrocyte density also required oligodendroglial expression of TrkB. It is unclear if this increase is due to TrkB stimulation of OPC proliferation, or enhanced survival. A role for BDNF in OPC proliferation has been suggested, with reduced OPC proliferation exhibited in BDNF deficient mice following cuprizone demyelination (VonDran *et al.*, 2011; Fulmer *et al.*, 2014). Despite reduced OPC proliferation, the density of mature oligodendrocytes was unchanged between BDNF deficient and wildtype mice following recovery (VonDran *et al.*, 2011). We have shown deletion of TrkB from maturing oligodendrocytes in development causes OPC hyperproliferation, but this too, exerts no effect on the density of mature oligodendrocytes in adulthood (Wong *et al.*, 2013). Collectively, these data suggest BDNF-TrkB signaling is not essential for oligodendrocyte differentiation, but rather that TrkB activation regulates OPC proliferation *in vivo*. Labelling proliferating cells at the time of cuprizone withdrawal and TDP6 delivery in wildtype and conditional knockout mice would determine whether TrkB signaling exerts a controlling influence upon OPC proliferation following myelin injury. Given the increase in Olig2^+^CC1^+^ oligodendrocytes in response to TDP6 depended on oligodendroglial expression of TrkB, sustained levels TrkB activation may also potentially override apoptotic signals. This would enhance OPC survival, subsequently increasing the density of differentiated oligodendrocytes. Regardless, the ultimate fate and persistence of all Olig2^+^CC1^+^ oligodendrocytes produced by TDP6 treatment in the long-term remyelinating CNS is an important question remaining to be answered, and is pertinent in the context of chronic demyelination in MS.

Intriguingly, while deletion of TrkB from oligodendrocytes abrogated the effects of TDP6 on myelin sheath thickness and oligodendrocyte differentiation, the proportion of myelinated axons still increased. This suggests that TDP6 may also have effects independent of oligodendroglial TrkB. The lack of a significant effect of TDP6 on astrogliosis and microgliosis suggests TDP6 does not directly modulate these cell populations, although potentially it alters the composition of their secreted factors (Djalali *et al.*, 2005). It is also possible increased myelination in the TDP6 treated conditional knockout mice was mediated by a small subset of oligodendrocytes that escape recombination and continue to express TrkB. This view would be consistent with previous work by us (Xiao *et al.*, 2011; Wong *et al.*, 2013) and others (Minichiello *et al.*, 1999; Du *et al.*, 2003; Medina *et al.*, 2004) indicating the role of TrkB in CNS myelination is specific to oligodendrocytes. However, what is striking is that the increase in myelinated axons in the conditional knockout mice treated with TDP6 was driven by an increase in the frequency of small dimeter axons (0.3-0.6μm) becoming myelinated. This is suggestive of a selective axonal response potentially driving initiation of myelination on this subpopulation of small caliber axons. The recent focus on activity-dependent myelination has indicated that efficient remyelination, on small diameter axons in particular, may require neuronal signals (Gautier *et al.*, 2015; Bechler, Swire and Ffrench-Constant, 2018). Optogenetic stimulation of cortical neurons increased neuronal secretion of BDNF (Venkatesh *et al.*, 2015), and this manipulation in healthy cortex resulted in increased callosal myelination (Gibson *et al.*, 2014). This suggests a possible role for neuronally derived autocrine BDNF-TrkB signaling in adaptive myelination, and importantly remyelination (Lundgaard *et al.*, 2013). This potential role for neuronally derived TrkB in promoting remyelination may be fortuitous since TrkB receptor expression is found in neurons adjacent to demyelinated white matter lesions in MS (Stadelmann *et al.*, 2002) and could be directly targeted with TDP6, or by other indirect strategies to sustain TrkB phosphorylation and remyelination. The mechanism mediating the TDP6-dependent increase in the frequency of myelinated axons in conditional knockout mice is unclear. Adoption of neuronal TrkB conditional knockout strategies will ultimately confirm whether neuronal TrkB directly influences remyelination, or whether this effect is due to an indirect response stimulated by TDP6.

We have demonstrated that TDP6, a structural mimetic of the TrkB binding region of BDNF, enhances remyelination in a preclinical animal model of MS. By targeting TrkB after demyelination, oligodendrocyte differentiation, the proportion of myelinated axons and myelin sheath thickness all increased. These are key outcomes for a potential remyelinating strategy. Deletion of TrkB from oligodendrocytes abrogated the effect of TDP6 on myelin sheath thickness and oligodendrocyte differentiation, but an increase in myelinated axons persisted, revealing a potential for targeting additional non-oligodendrocyte specific effects of TrkB in remyelination. Overall, our results support the development of therapeutic strategies to sustain elevations of TrkB signaling to promote myelin repair in demyelinating disease.

## Materials and methods

### Experimental animals and cuprizone-induced demyelination

Female C57BL/6 mice, aged 8-10 weeks were fed 0.2% cuprizone in normal chow (Teklad Custom Research Diets, USA) for 6 weeks to induce demyelination. Cuprizone feed was removed, and mice were sacrificed or received intracerebroventricular osmotic pumps for 7 days.

For experiments in conditional knockout mice, 8-10 week old CNPaseCre^+/−^ female progeny of CNPaseCre^+/−^ mice (Lappe-Siefke *et al.*, 2003) crossed to TrkB^fl/fl^ mice (Lulkart *et al.*, 2005) bred to a C57BL/6 background for five generations, underwent procedures described above. Additional Cre^+/−^ and Cre^+/−^ female mice aged 14-16 weeks were used as healthy controls.

All mice were housed in specific pathogen-free conditions at the Melbourne Brain Centre Animal Facility. All animal procedures were approved by The Florey Institute for Neuroscience and Mental Health Animal Ethics Committee and followed the Australian Code of Practice for the Care and Use of Animals for Scientific Purposes.

### Intracerebroventricular delivery of BDNF and TDP6

Following cuprizone feeding, mice received either 44μM BDNF carried in 0.1% BSA in artificial CSF (aCSF, *n*=8), 40μM TDP6 in aCSF (n=9) or the aCSF vehicle (n=7) via intracerebroventricular osmotic pumps with a flow rate of 0.5 uL/hr (Azlet, CA, USA). Infusion concentrations were based on effective concentrations of BDNF and TDP6 in previously published *in vitro* myelination assays [13]. Cannulae were implanted at coordinates +0.5mm rostral and −0.7mm lateral of Bregma to administer into the right lateral ventricle. Anesthesia was induced with 4-5% isoflurane and 0.5% oxygen, and maintained at 2.5-1% isoflurane and 0.5% oxygen through a nose cone during stereotaxic manipulation. All mice were placed in a recovery chamber maintained at 32°C and monitored immediately following surgery for adverse reactions, and then daily. After 7 days of continuous infusion, mice were sacrificed and the brain removed for immunostaining and electron microscopic (EM)analysis.

### Tissue processing and immunofluorescence

Mice were anaesthetized and transcardiallly perfused with 0.1M sterile mouse isotonic phosphate buffered saline (PBS) followed by 4% paraformaldehyde (PFA). Brains were collected and post-fixed in 4% PFA overnight. The first millimeter of the right hemisphere from the sagittal midline was selected using a sagittal mouse brain matrix and placed in Kanovsky’s buffer overnight, and washed in 0.1M sodium cacodylate before embedding in epoxy resin at the Peter MacCallum Centre for Advanced Histology and Microscopy for EM analysis. The left hemisphere was cryoprotected in 30% sucrose and frozen in OCT in isopentane over dry ice. Sagittal sections were cut at 10μm using a cryostat maintained between −20 to −17°C and collected on Superfrost+ slides, air-dried and stored at −80°C until use. Approximately 70-100μm separated adjacent sections on each slide. Sections cut beyond ±2.64mm lateral from the midline were excluded.

For immunofluorescence, slides were washed in PBS prior to overnight incubation at room temperature with primary antibodies diluted in 10% normal donkey serum (NDS) with 0.3% Triton-X 100. Slides were washed in PBS before 2-hours incubation with the appropriate fluorophore-conjugated secondary antibody in the dark. After washing with PBS, slides were counterstained with nuclear marker Hoechst 33442 and cover slipped using aqueous mounting media (Dako, CA, USA). For all stains, immunohistochemistry was performed in batches.

Antibodies used were: rat anti-myelin basic protein (MBP, 1:200, MAB386, Millipore, MA, USA) as a marker for remyelination, rabbit anti-Olig2 (1:200, AB9610, Millipore, MA, USA), mouse anti-CC1 (1:200, APC, OP80, CalBioChem, CA, USA), goat anti-platelet derived growth factor receptor-alpha (PDGFRα, 1:200, AF1062, R&D Systems, MN, USA) to identify stages of the oligodendrocyte lineage, goat anti-Iba1 (Iba1, 1:200, ab5076, Abcam, UK) for microglia and mouse anti-glial fibrillary acidic protein (GFAP, 1:100, MA360, Millipore, MA, USA) for astrocytes. To identify the level of TrkB expression rabbit anti-TrkB (1:500, R-149-100, Biosensis, SA, Australia) was used, and TrkB activation was detected with antibodies against phosphorylated TrkB (pTrkB^S478^, 1:200, R-1718-50, Biosensis).

For MBP immunostaining sections were post-fixed with ice-cold 100% methanol for 10mins prior to the first wash. For pTrkB^S478^, tris-based saline (TBS) with 0.3% Triton-X 100 was used for all washes and the antibody diluent contained 1% bovine serum albumin in addition to 10% NDS and primary antibody incubation was performed overnight at 4°C.

### Electron microscopy and analysis

Semi-thin (0.5-1μm) sections of the caudal corpus callosum in a sagittal plane were collected on glass slides and stained with 1% toluidine blue. Ultrathin (0.1μm) sections were subsequently collected on 3⨯3mm copper grids. Specimens were examined using JEOL 1011 transmission electron microscope, and images were collected using MegaView III CCD cooled camera operated with iTEM AnalySIS software (Olympus Soft Imaging Systems GmbH). Six distinct fields of view were imaged at 10000x magnification per animal. Images were used to count myelinated axons, measure axon diameters, myelin thickness and g-ratios in FIJI/ImageJ (ImageJ 1.51K, National Institutes of Health). A minimum of three fields of view (142μm^2^) were examined per animal with 3-4 mice per treatment group. For g-ratios at least 100 axons from 3 mice per group were measured. Resin embedding, sectioning, post-staining and EM imaging were performed at the Peter MacCallum Centre for Advanced Histology and Microscopy.

### Fluorescence imaging and analysis

All imaging was performed blinded to treatment group and restricted to the corpus callosum, approximately -1.1mm to -3.0mm from Bregma. For each analysis, a minimum of three sections per animal were imaged.

To quantify the level of remyelination images of MBP-stained sections were collected with an AxioVision Hr camera attached to a Zeiss Axioplan2 epi-fluorescence microscope under a 20x objective. Uniform exposure times were used.

Remaining analyses were performed using images acquired with a Zeiss LSM780 or LSM880 confocal microscope with 405nm, 488nm, 561nm and 633nm laser lines.
For each fluorescent stain uniform settings were used.

MBP and GFAP staining were quantified as described in (Fletcher *et al.*, 2014) using the threshold function in FIJI/ImageJ and limited to a standard region of interest (ROI) of 625000μm^2^ for each section. Data were expressed as the percentage area of positive staining in a single ROI.

### Cell counts

All cell counts were performed using maximum intensity projection images generated from z-stacks. A standard ROI was set (625000μm^2^) and nuclei masks identified by Hoechst or positive Olig2 staining were segmented and counted in FIJI/Image J using the “Analyze Particles…” function to create masks. The threshold function was then applied to identified positive CC1, PDGFRα, Iba1 positive staining and to generate a binary image. Positive nuclei were identified using the Binary Reconstruction plugin (Legland, Arganda-Carreras and Andrey, 2016), and counted with the Analyze Particles function. Automated counts were verified by manually counting a sub-sample of images. For pTrkB^S478^ cell counts were performed manually. Data were expressed as the number of cells/mm^2^ or the proportion out of the total number of nuclei.

### High performance liquid chromatography and mass spectroscopy analysis of TDP6 after 7 days mini-pump incubation *in vivo*

TDP6 prepared the day prior to pump administration (day 0) and retrieved from the reservoir of osmotic mini-pumps implanted in conditional knockout mice (day 7) were analyzed by reverse-phase high performance liquid chromatograph (RP-HPLC), using an Agilent 1200 series unit, fitted with a Phenomenex Luna C8 column (5u; 50⨯4.6mm), running 0-60% acetonitrile over a 14-min window. UV spectra were measured for each sample using a 214nm wavelength, showing a single TDP6 peptide peak at 7.3min (Fig. S3). A sample of aCSF was run as a control. Liquid-chromatography mass spectrometry (LCMS), using an Agilent 6100 series single quadrupole system was used to confirm the molecular weight of the peptide, TDP6 showing the predicted *m/z* of 1207.5 [M + 2H]^2+^.

### Statistical analyses

All data collection was performed blind to sample identity. To compare between treatment groups, 1-way ANOVA or unpaired Student’s t-tests were performed with post-hoc multiple comparison tests as appropriate. For g-ratios, linear regressions were performed, and axon diameter frequency distribution were assessed using χ^2^ distribution tests. All statistical tests were performed in GraphPad Prism 7 (GraphPad Software Inc.) with p<0.05 considered significant.

## Acknowledgments

This work was supported by an Australian National Health and Medical Research Council (NHMRC) Project Grants (APP1058647 and APP1105108; SSM and JX) and a Multiple Sclerosis Research Australia (MSRA) Project Grant (13-039; SSM and JX). JLF was supported by a MSRA Postdoctoral Fellowship (14-056). DGG was supported by an MSRA/NHMRC Early Career Fellowship (APP111041). We thank Cameron Nowell (Monash University) and Dr Verena Wimmer (The Florey Institute) for advice on image analysis procedures and automation. Confocal imaging was performed using facilities provided by the Biological Optical Microscope Platform, The University of Melbourne, and the Florey Advanced Microscopy and Immunohistochemistry Service, The Florey Institute of Mental Health and Neuroscience. We acknowledge the Peter MacCallum Centre for Advanced Histology and Microscopy for assistance with EM processing and imaging.

## Supplementary Information

**Fig. S1:**
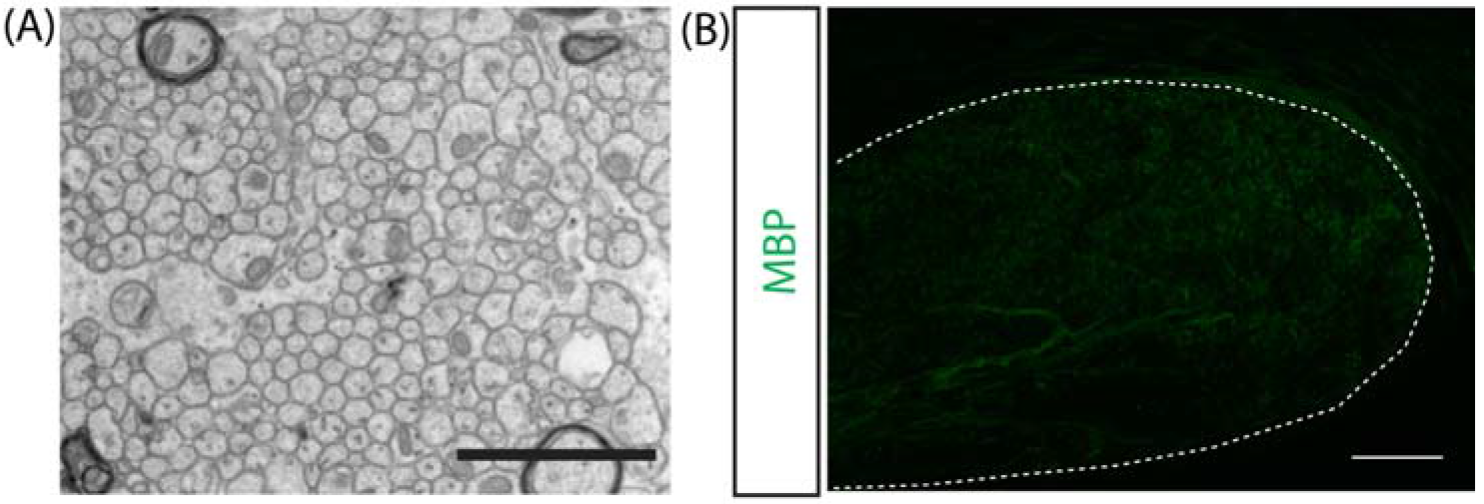
Demyelination in the caudal corpus callosum of C57BL/6 mice fed 0.2% cuprizone for 6 weeks. **(A)** Representative electron micrograph of demyelinated axons in the caudal corpus callosum of a 14-week old mouse. Scale bar=2μm. **(B)** Representative micrograph of myelin basic protein (MBP) immunostaining in the caudal corpus callosum (outlined). Scale bar=100μm.

**Fig. S2:**
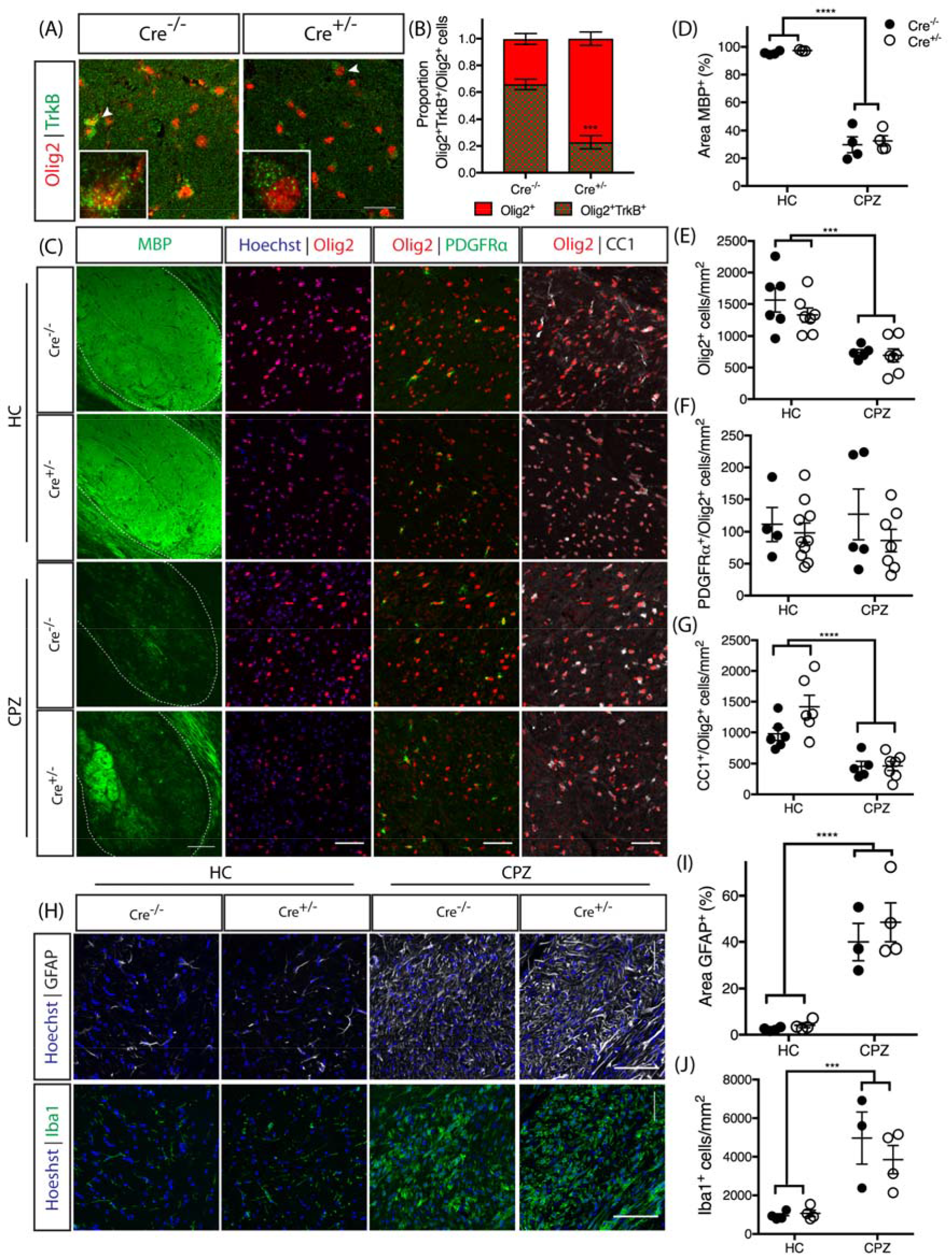
CNPaseCre^+/−^ x TrkB^fl/fl^ mice exhibit normal adult myelination and response to cuprizone-demyelination. **(A)** Representative micrographs of TrkB and Olig2 immunostaining in the caudal corpus callosum of CNPaseCre^+/−^ and CNPaseCre^+/−^ TrkB^fl/fl^ healthy controls aged 16 weeks. Scale bar=20μm **(B)** Proportion of Olig2^+^ cells positive for TrkB was significantly decreased in CNPaseCre^+/−^ TrkB^fl/fl^ mice compared to Cre^+/−^ controls. Student’s t-test, *n*=4-6/group,***p=0.0003. Mean ± SEM plotted. **(C)** Representative micrographs of immunostaining for MBP and oligodendrocyte lineage markers Olig2, PDGFRP, and CC1 in the caudal corpus callosum of CNPaseCre^+/−^ (closed circles) and CNPaseCre^+/−^ (open circles) TrkB^fl/fl^ healthy controls (HC) and cuprizone-fed (CPZ) mice. Scale bar=50μm. **(D)** Area of MBP immunostaining and **(E)** Olig2^+^ cell density were unchanged due to genotype in the healthy controls, but were significantly decreased in both genotypes following cuprizone treatment (****p<0.0001 and ****p<0.0001, respectively). **(F)** Density of Olig2^+^PDGFRαα^+^ OPCs was unchanged with cuprizone treatment or genotype (p=0.94). **(G)** Olig2^+^CC1^+^ oligodendrocytes were unchanged between genotypes in healthy controls, while cuprizone feeding significantly reduced the density of these cells in the caudal corpus callosum of both genotypes (****p<0.0001). **(H)** Representative micrographs of GFAPα^+^ astrocytosis and Iba1^+^ microglia in the caudal corpuscallosum of Cre^+/−^ and Cre^+/−^ HC andCPZ mice. Scale bar=50μm. **(I)** Percentage area GFAP^+^ and **(J)** density of Iba1^+^ microglia were similar in both genotypes in HC, and were increased in both genotypes following CPZ (****p<0.0001 and ***p=0.0003). For *(D-G)* and *(I-J*), *n*=3-7/group, 2-way ANOVA.

**Fig. S3:**
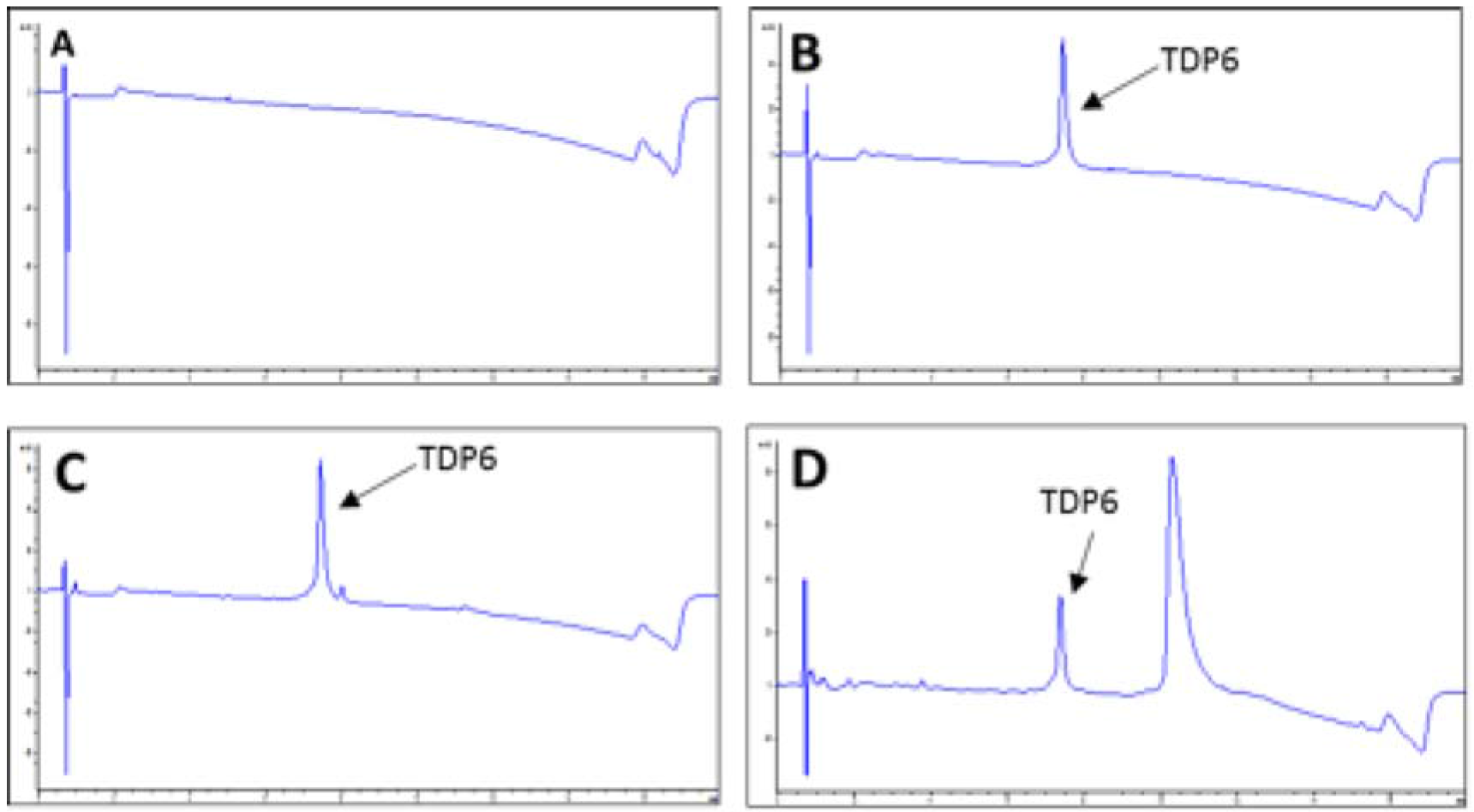
TDP6 was retrieved and detected after 7-days incubation within osmotic mini-pumps implanted in conditional knockout mice. **(A)** Reverse-phase high-performance liquid chromatography (HPLC) UV trace of artificial CSF(‘blank/control’ sample).**(B)** HPLC UV trace of TDP6 at day prior to animal administration. **(C)** HPLC UV trace of TDP6 at day 0. (**D)** HPLC UV trace of TDP6 at day 7. All traces measured using 214nm wavelength. **Note:** Additional peak in (D) was determined to not be a peptide degradation product. Area under the peptide peak was quantified in (B-D) and no significant change was observed.

